# Chicago and Dovetail Hi-C proximity ligation yield chromosome length scaffolds of *Ixodes scapularis* genome

**DOI:** 10.1101/392126

**Authors:** Andrew B. Nuss, Arvind Sharma, Monika Gulia-Nuss

## Abstract

A high-quality genome sequence is essential for understanding an organism on molecular level. However, the larger genomes with substantial repetitive sequences are challenging to assemble with the sequencing technologies. Hi-C technique is changing the genome architecture landscape by providing links across a variety of length scales, spanning even whole chromosomes. *Ixodes scapularis* haploid genome is 2.1 gbp and the current assembly consists of 369,495 scaffolds representing 57% of the genome. The fragmented genome poses challenges with functional gene analysis and an improved assembly is needed. We therefore used the Hi C technique to achieve chromosomal level assembly of tick genome. With Chicago and Dovetail Hi C assemblies, we were able to achieve 28 >10Mb sequences that correspond to 28 chromosomes in *I. scapularis*.

## Introduction

*Ixodes scapularis* is the principal vector of the Lyme disease spirochete, *Borrelia burgdorferi*. *I. scapularis* genome was the first and only medically important acarine species sequenced and annotated thus far. The genome was sequenced using BAC clones and Sanger sequencing methods. However, the 2.1 gb haploid genome with long repetitive sequencing posed challenge to achieve scaffolds that span entire chromosomes. Some repetitive regions were too large and difficult to be spanned by the available clone libraries. The assembly, IscaW1, comprises 369,495 scaffolds representing 57% of the genome. The fragmented genome further poses challenges in identifying gene sequences and therefore a high quality genome sequence is needed for advance genomics and genetics work in this vector.

A high-quality genome sequence is essential for understanding an organism on molecular level. However, the larger genomes that contain substantial repetitive sequences as the case of *I. scapularis* genome are challenging to assemble with the current technologies.

The availability of sequencing methods that could produce scaffold size sequence length and three dimensional chromatin capture such as PacBio, and HiC, respectively, are changing the genome sequencing landscape. Hi-C is a sequencing-based approach for determining how a genome is folded by measuring the frequency of contact between pairs of loci (Lieberman-Aiden eta l., 2009; Rao et al., 2014). Contact frequency depends strongly on the base pair distance between a pair of loci. Hi-C data can provide links across a variety of length scales, spanning even whole chromosomes. Hi-C has been used to improve draft genome assemblies (Marie-Nelly et al., 2014; Dudchenko et al., 2017) and to create chromosome-length scaffolds for large genomes (Burton et al., 2013; Dudchenko et al., 2017). We therefore used the Hi C technique to achieve chromosomal level assembly of tick genome. We carried out Chicago and Hi C assemblies that utilize *in vitro* and *in vivo* chromatin structures, respectively, in order to provide the best scaffolding success. We successfully assembled the genome in 28 >10Mb sequences that correspond to 28 chromosomes in *I. scapularis*.

## Methods

### Sample collection

Fully engorged *I. scapularis* females were obtained from the Oklahoma State University and maintained in our colony at 20°C and 98% relative humidity. Three batches of eggs (approximately 3000 eggs each) from three individual females were flash frozen in liquid nitrogen and shipped to Dovetail genomics for Chicago and HiC assemblies.

### Chicago library preparation and sequencing

Three Chicago libraries were prepared as described previously (Putnam et al, 2016). Briefly, for each library, ~500ng of high molecular weight genomic DNA (gDNA) with mean fragment length = 100 was reconstituted into chromatin *in vitro* and fixed with formaldehyde. Fixed chromatin was digested with DpnII, the 5’ overhangs filled in with biotinylated nucleotides, and then free blunt ends were ligated. After ligation, crosslinks were reversed and the DNA purified from protein. Purified DNA was treated to remove biotin that was not internal to ligated fragments. The DNA was then sheared to ~350 bp mean fragment size and sequencing libraries were generated using NEB Next Ultra enzymes and Illumina-compatible adapters. Biotin-containing fragments were isolated using streptavidin beads before PCR enrichment of each library. The libraries were sequenced on an Illumina HiSeq X (rapid run mode).

### Hi C library preparation and sequencing

Three HiC libraries were prepared in a similar manner as described previously (Erez Lieberman-Aiden et al., 2009). Briefly, for each library, chromatin was fixed in place with formaldehyde in the nucleus and then extracted. Fixed chromatin was digested with DpnII, the 5’ overhangs filled in with biotinylated nucleotides, and then free blunt ends were ligated. After ligation, crosslinks were reversed and the DNA purified from protein. Purified DNA was treated to remove biotin that was not internal to ligated fragments. DNA was then sheared to ~350 bp mean fragment size and sequencing libraries were generated using NEB Next Ultra enzymes and Illumina-compatible adapters. Biotin-containing fragments were isolated using streptavidin beads before PCR enrichment of each library. The libraries were sequenced on an Illumina HiSeq X (rapid run mode).

### Scaffolding the assembly with HiRise

The input *de novo* assembly, shotgun reads, Chicago library reads, and Dovetail HiC library reads were used as input data for HiRise, a software pipeline designed specifically for using proximity ligation data to scaffold genome assemblies (Putnam et al, 2016). An iterative analysis was conducted. First, Shotgun and Chicago library sequences were aligned to the draft input assembly using a modified SNAP read mapper (http://snap.cs.berkeley.edu). The separations of Chicago read pairs mapped within draft scaffolds were analyzed by HiRise to produce a likelihood model for genomic distance between read pairs, and the model was used to identify and break putative misjoins, to score prospective joins, and make joins above a threshold. After aligning and scaffolding Chicago data, Dovetail HiC library sequences were aligned and scaffolded following the same method. After scaffolding, shotgun sequences were used to close gaps between contigs (Fig. 1).

**Figure 1:**
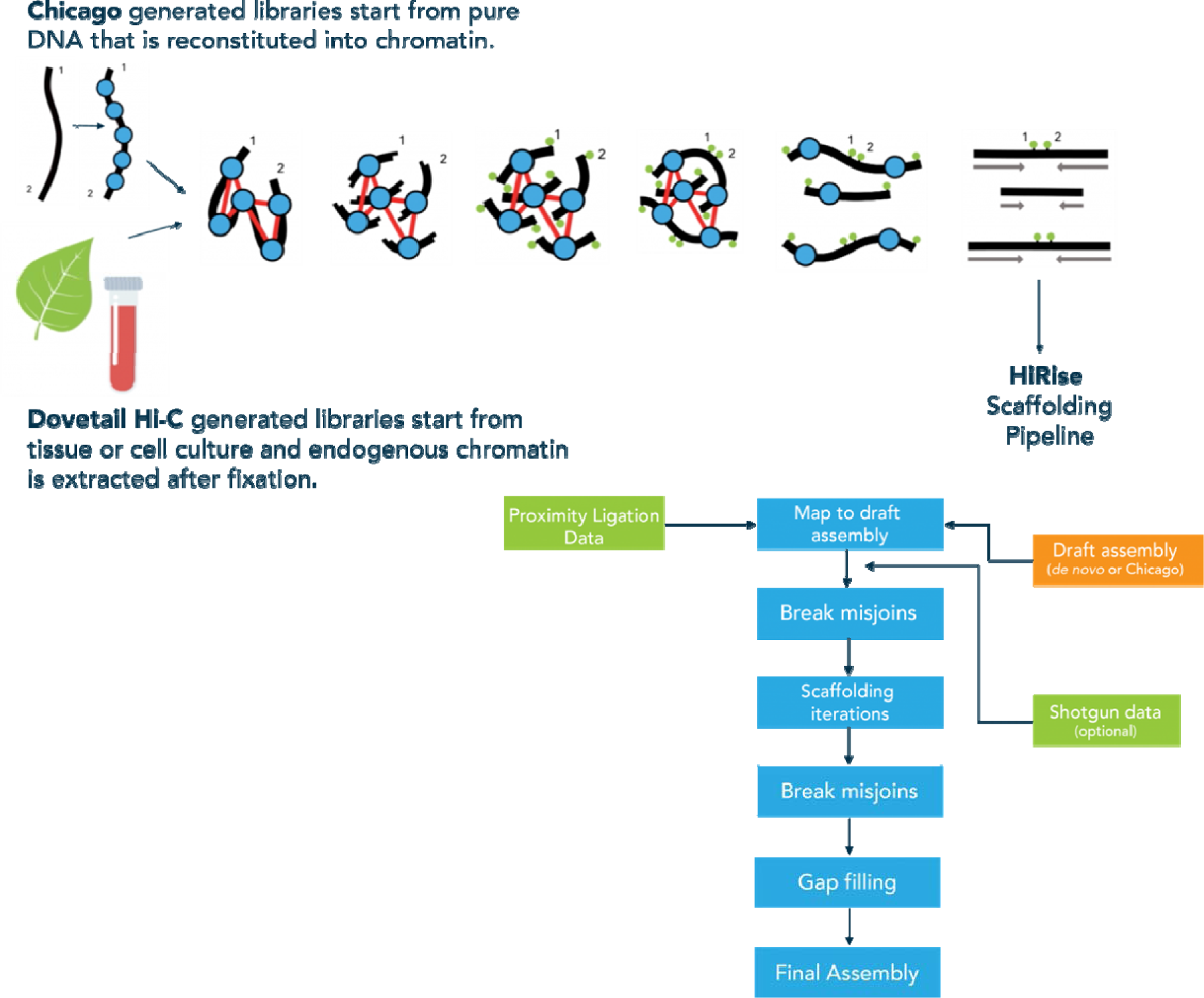
Schematic of Chicago and Dovetail library preparation and Final HiRise scaffolding pipeline.

## Results and Discussion

For Chicago libraries: each library produced 153 million reads of 2 × 151 bp length sequences. Together, these Chicago library reads provided 25.72 x physical coverage of the genome (1-100 kb pairs).

For Hi C libraries: Library 1 produced 143 million; library 2 produced 162 million; and library 3 produced 123 million reads of 2 × 151 bp length. Together, these HiC library reads provided 686.76 x physical coverage of the genome (10-10,000 kb pairs).

With Chicago and HiC libraries assembled with HiRise, we were able to significantly improve the existing IscaW1 assembly. Chicago assembly resulted in L50/N50 of 1,031 scaffolds and L90/N90 of 6,599 scaffolds (Table- 1). Using Chicago (*in vitro*) as an input assembly we performed HiRise assembly with the sequences obtained from HiC sequencing (*in situ*). The HiRise assembly resulted in L50/N50 of 12 scaffolds that were >48 Mb and 28 scaffolds at >10 Mb (Table-1). It significantly improve L90/N90 of our input (Chicago) assembly and resulted in 2,895 scaffolds. A comparison of the contiguity of the input assembly and the final HiRise scaffolds curve also showed the increased scaffold length suggesting a much improved assembly (Fig. 2). Scaffolds less than 1 kb were excluded from the analysis. Estimated physical coverage (10-10,000 kb pairs) was 686.76X (Fig. 3).

**Figure 2:**
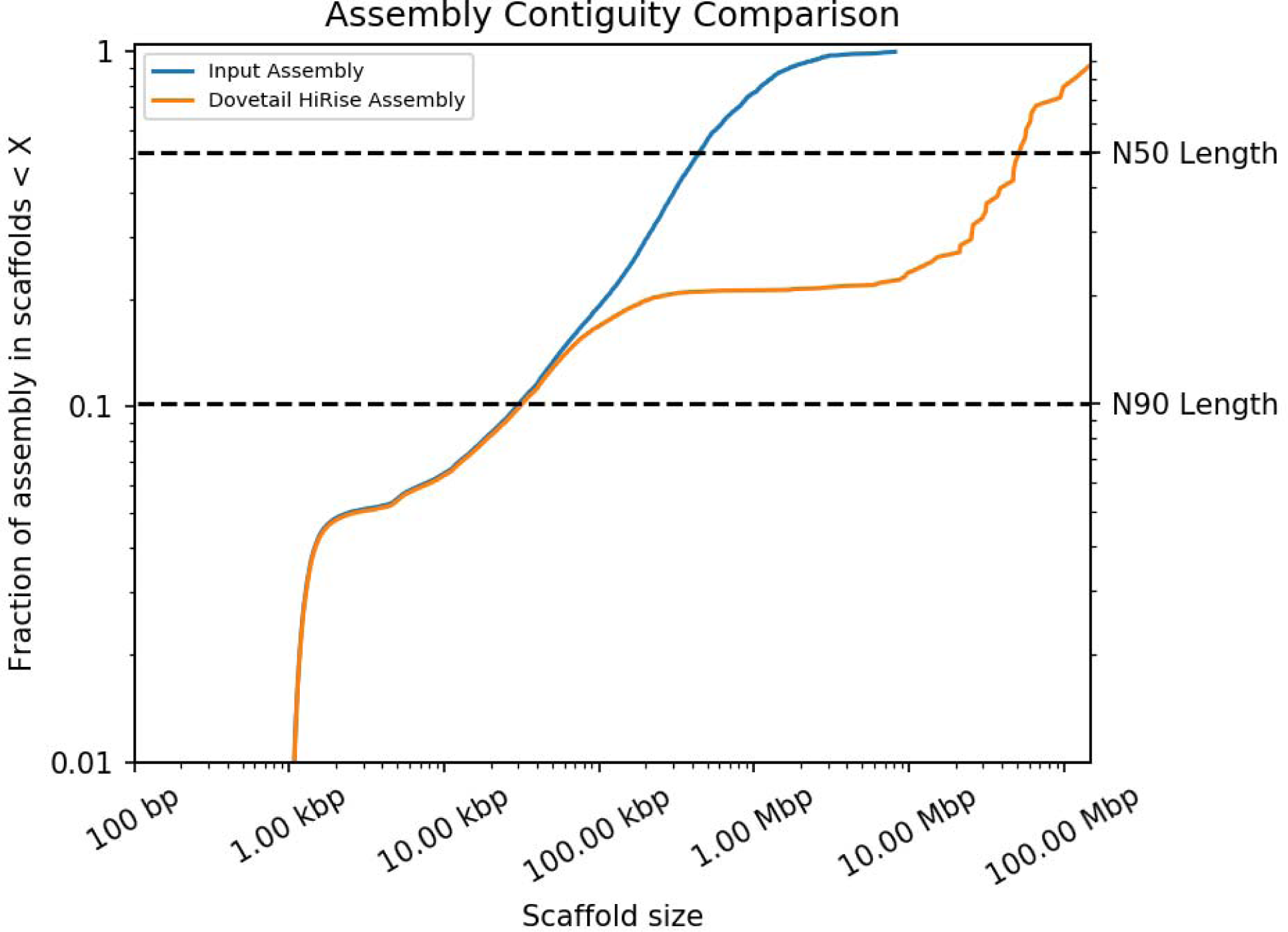
A comparison of the contiguity of the input assembly and the final HiRise scaffolds. Each curve shows the fraction of the total length of the assembly present in scaffolds of a given length or smaller. The fraction of the assembly is indicated on the Y-axis and the scaffold length in basepairs is given on the X-axis. The two dashed lines mark the N50 and N90 lengths of each assembly. Scaffolds less than 1 kb are excluded.

**Figure 3:**
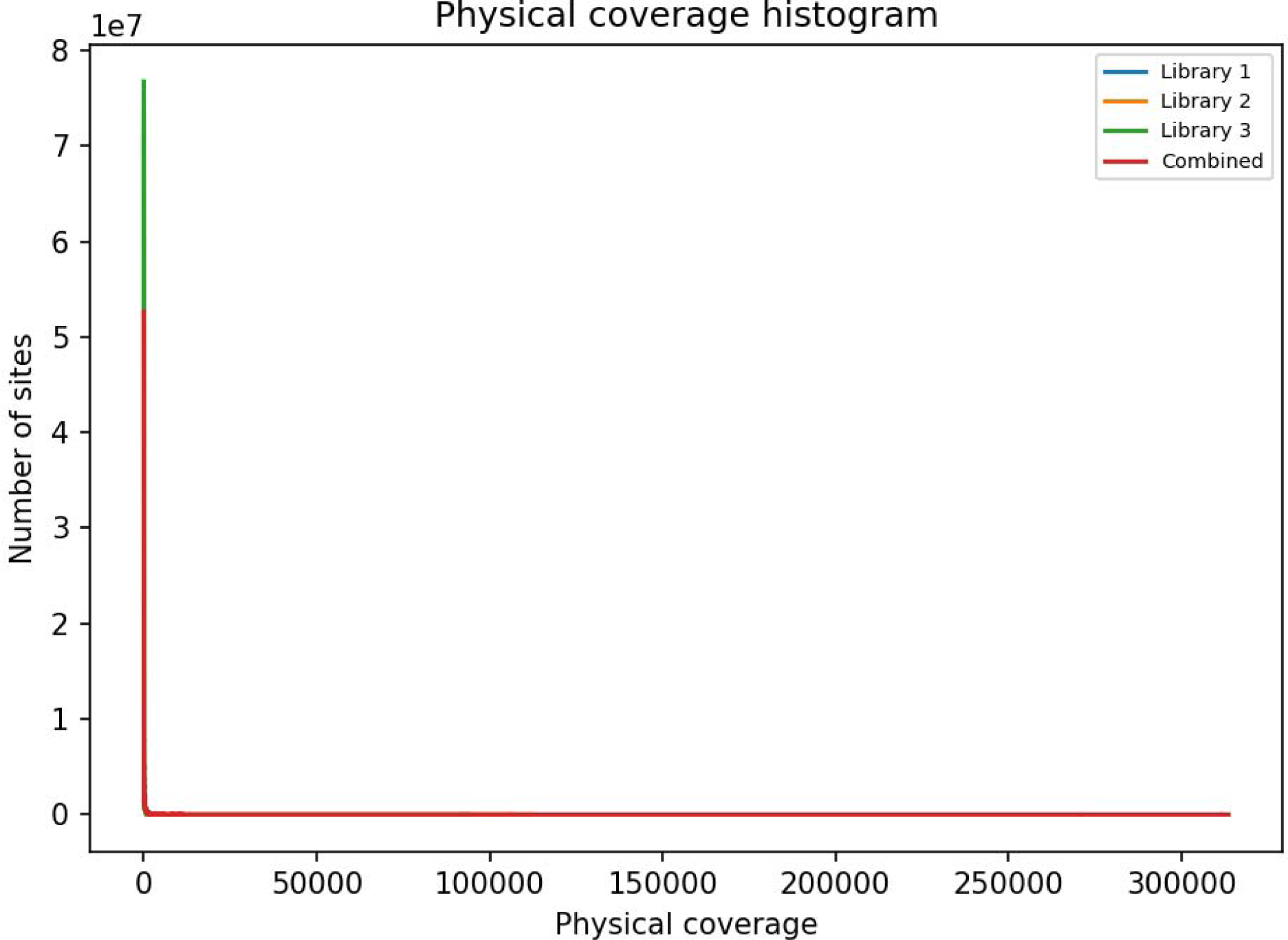
Histogram of physical coverage over input assembly. Coverage values are calculated as the number of read pairs with inserts between 10 and 10,000 kb spanning each position in the input assembly.

We used a robust procedure for Hi-C linking information to generate accurate genome assemblies with chromosome-length scaffolds. We first used Chicago data to identify and correct errors in the scaffolds of the initial Iscaw1 assembly by correcting misjoins. Next, we used the Chicago assembly data as our input assembly and corrected misjoins with Hi C data. We used a novel HiRise algorithm to anchor, order, and orient the resulting sequences, employing the contact frequency between a pair of sequences as an indicator of their proximity in the genome. Finally, we merged contigs and scaffolds that correspond to overlapping regions of the genome by identifying pairs of scaffolds that exhibit both strong sequence homology as well as strong similarity in long-range contact pattern (Fig. 4). Scaffolds less than 1 Mb were excluded from the analysis.

**Figure 4:**
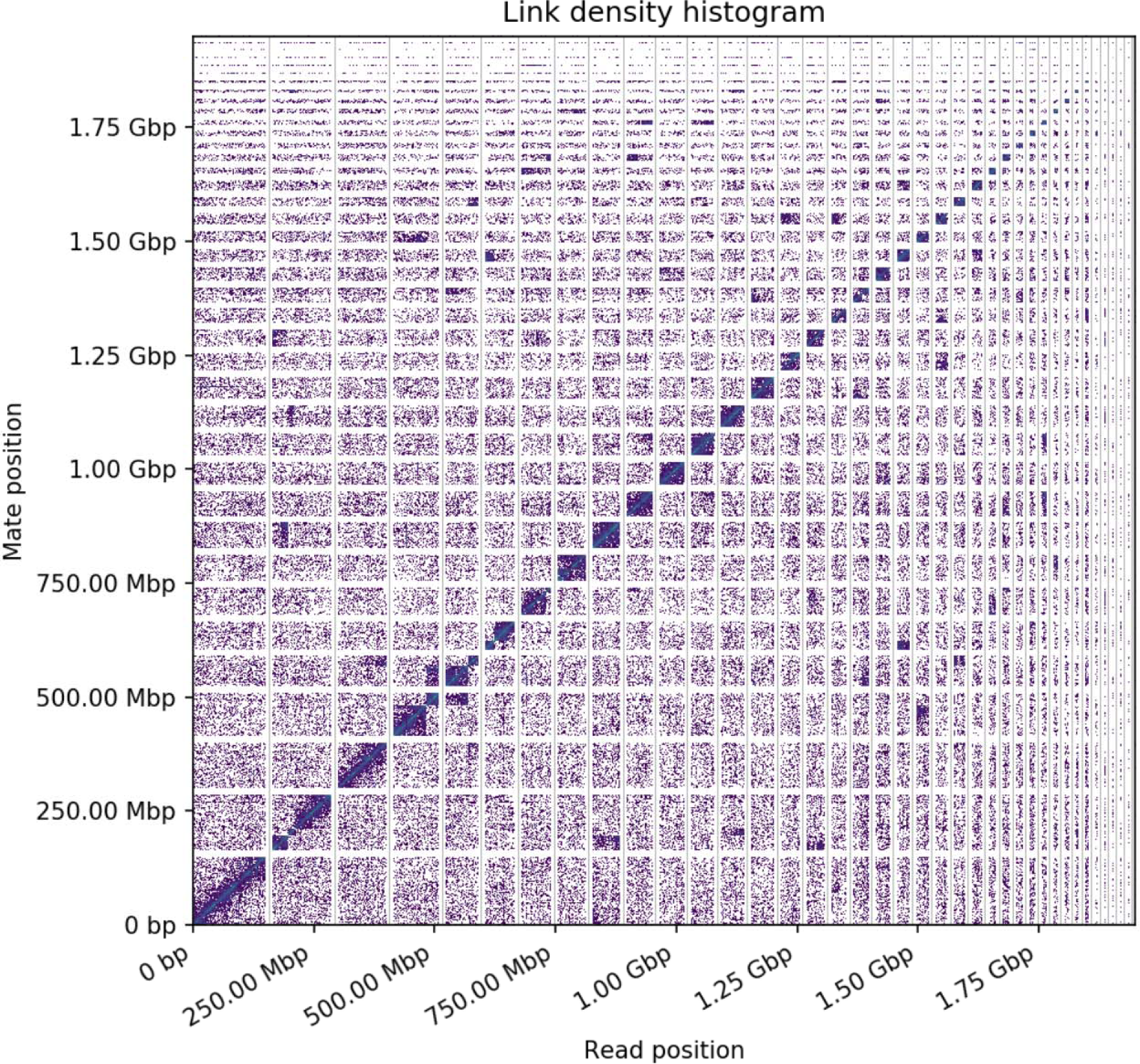
Link density histogram. The x and y axes give the mapping positions of the first and second read in the read pair respectively, grouped into bins. The color of each square gives the number of read pairs within that bin. White vertical and black horizontal lines have been added to show the borders between scaffolds. Scaffolds less than 1 Mb are excluded.

After Chicago and HiC assemblies with HiRise, we were able to obtain a longest scaffold of 1.49 Gb. The final assembly resulted in 83, 347 scaffolds and contig N50 of

64.21 Kb compared with 369, 496 scaffolds and contig N50 of 2.94 Kb of the original IscaW genome. There was 2.85 % genome still in gaps but that is a significant decrease from ~49% genome in gaps from the original IscaW assembly (Gulia-Nuss et al., 2016)

Overall, the HiRise assembly (Chicago+ HiC) resulted in 28 scaffolds longer than 10 Mb. This number corresponds to the chromosome number in *I. scapularis* genome (Gulia-Nuss et al., 2016) suggesting a chromosome level assembly. Our results show that Hi-C data provides a rapid, inexpensive method for generating de novo assemblies with chromosome-length scaffolds. It is important to bear in mind that these assemblies still contain gaps. Additional data such as PacBio and 10X sequences could improve the results. We are now planning to fill the gaps in this assembly with PacBio sequencing and redo the scaffolding based on the additional data. The high contiguity of the data is a marked improvement over existing assembly. We understand that mapping all of the existing gene models onto these scaffolds will be challenging, however, once done this will be allow sophisticated genomics and genetics research in this disease vector that is currently not possible.

In conclusion, the ability to rapidly and reliably generate genome assemblies with chromosome-length scaffolds for non-model organisms should accelerate genomic analysis of many organisms.

## Acknowledgements

This project was funded through a matching grant GUL355 by the Dovetail Genomics. We thank Mark Daly and Lily Shiue from Dovetail Genomics for library preparations, sequencing, and data analysis. We also thank Dr. Won Choel Yim for insightful discussions.

---

**Table 1:** L50/ L90 lengths after Chicago and Doveatil HiC assemblies.

|  | **Input Assembly** | **Dovetail HiRise Assembly** |
| --- | --- | --- |
| Total length | 1,794.02 Mb | 1,794.46 Mb |
| L50/N50 | 1,031 scaffolds; 0.419 Mb | 12 scaffolds; 48.591 Mb |
| L90/N90 | 6,599 scaffolds; 0.029 Mb | 2,895 scaffolds; 0.030 Mb |

**Table 2:** Comparative statistics of new assemblies with existing IscaW assembly. N50 is defined as the scaffold length such that the sum of the lengths of all scaffolds of this size or less is equal to 50% of the total assembly length

| **Comparative Assembly Statistics** |  |  |  |
| --- | --- | --- | --- |
|  | **Input Assembly** | **HiRise Assembly** | **Original IscaW Assembly** |
| Longest Scaffold | 8,139,475 bp | 149,656,909 bp | - |
| No. of Scaffolds | 87,681 | 83,347 | 369,495 |
| No. of scaffolds >1Kb | 87,006 | 82,672 | ~ 369,000 |
| Contig N50 | 64.29 kb | 64.21 kb | 2.94 kb |
| No. of gaps | 286,616 | 291,041 | - |
| Percent genome in gaps | 2.83% | 2.85% | ~49% |

